# Genetic mixing and demixing on expanding spherical frontiers

**DOI:** 10.1101/2023.07.07.548058

**Authors:** Alba García Vázquez, Namiko Mitarai, Liselotte Jauffred

## Abstract

Genetic fluctuation during range expansion is a key process driving evolution. When a bacterial population is expanding on a 2D surface, random fluctuations in the growth of the pioneers at the front line cause a strong de-mixing of genotypes. Even when there is no selective advantage, sectors of low genetic diversity are formed. Experimental studies of range expansions in surface-attached colonies of fluorescently-labeled microorganisms have contributed significantly to our understanding of fundamental evolutionary dynamics. However, experimental studies on genetic fluctuations in 3D range expansions have been sparse, despite their importance for tumour or biofilm development. We encapsulated populations of two fluorescent *Escherichia coli* strains in inoculation droplets (volumes ∼0.1 nl). The confined ensemble of cells grew when embedded in a hydrogel – with nutrients – and developed 3D colonies with well-defined, sector-like regions. Using a confocal laser scanning microscope (CLSM), we imaged the development of 3D colonies and the emergence of sectors. We characterised how cell concentration in the inoculation droplet controls sectors, growth rate, and the transition from branched colonies to quasi-spherical colonies. We further analysed how sectors on the surface change over time. We complement these experimental results with a modified 3D Eden growth model. The model in 3D spherical growth predicts a phase, where sectors are merging, followed by a steady increase (constant rate), and the experimentally analysed sectors were consistent with this prediction. Ergo, our results demonstrate qualitative differences between radial (2D) and spherical (3D) range expansions and their importance in gene fixation processes.

## INTRODUCTION

In nature, bacteria and other single cellular organisms live in communities and often form dense structured populations such as colonies or biofilms. The structure, function, and stability of these communities depend on a complex network of social interactions, where bacteria exchange signals and metabolites, and protect each other from toxins, while at the same time proliferating and competing for space. This continuous cooperation and competition in the communities lead to complex spatial structures.

The structure has been widely investigated in surface-colonising microbial populations (see Ref. Eigentler et al. 2022a for a thorough review). These so-called *competition experiments*, in which well-mixed populations of bacteria (or yeast) Hallatschek and Nelson 2010 are inoculated on agar-surfaces and incubated, allow the cells to grow and divide and ultimately form complex macroscopic patterns Branda et al. 2005. When the initial founder cells are a mixture of differently colored fluorescent cells, one can observe patterns of spatially segregated lineages, even among bacteria of similar (i.e. neutral) fitness. It is well-known that the emergence of these sector-like regions is driven by random fluctuations at the outermost band of the expanding frontier Hallatschek et al. 2007. Also that the sector boundaries are diffusive Hallatschek and Nelson 2010, Korolev et al. 2011 and depend on, e.g., environmental conditions Excoffier and Ray 2008, Jauffred et al. 2017, extracellular matrix production Booth and Rice 2020, and cell shape Smith et al. 2017. Furthermore, the initial inoculating concentration has been found to control segregation patterns. In particular, the average sector regions’ sizes correlate with inoculation density van Gestel et al. 2014, Blanchard and Lu 2015, Borenstein et al. 2015, Bottery et al. 2019, Lee et al. 2022, as space limits proliferation during range expansion Eigentler et al. 2022b. These observations provide us with a deeper understanding of the population structure in biofilm and also serve as the foundation to understand the genetic drift and fixation in expanding populations in 2D Hallatschek et al. 2007, Korolev et al. 2010, Fusco et al. 2016.

Bacteria also live in 3D habitats, often in dense environments, where they are mired in mucus or entangled in polymers secreted by other bacteria, algae, or animal tissue. One model system of bacteria living in such settings is monoclonal spherical colonies in hydrogel Jung et al. 2015. The system has been used to investigate, e.g., growth Strathmann et al. 2000, Burmølle et al. 2009, colony morphology Zhang et al. 2021, Cordero et al. 2023, quorum sensing Müller et al. 2022, and phage sensitivity Eriksen et al. 2018.

Despite these advances in 3D model systems and the broad interest in competition and evolution in microbial communities as well as tumor growth Fusco et al. 2016, Ben-Jacob et al. 2012, we still lack a suitable experimental model system to study competition in growing 3D bacterial colonies. Here, we propose such a system: We used a 1:1 co-culture of two fluorescently labeled *E. coli* strains, which we encapsulated in agarose beads. We submerged these *inoculation beads* in a semi-solid agar (0.5%) and incubated them for a 3D competition assay. We imaged the resulting colonies using a confocal laser-scanning microscope (CLSM), to know how the density of founder cells impacts growth and pattern-formation. We further analyzed how the number of sectors on the colony surface changes with time. Our findings are complemented with mathematical modeling to provide insight into spatial segregation in 3D communities. In particular, we demonstrate that the sector splitting, which we also observe in our experiment, becomes dominant in the long term. Our system serves as an effective tool to investigate the multi-species bacterial community and also as an ideal model system to analyze spatial population genetics in 3D growing colonies.

## RESULTS

We mixed two sub-populations of the non-motile *E. coli* B strain (REL) carrying plasmids coding for either green fluorescent protein (GFP) or red fluorescent protein (RFP) in 1:1 ratio, as shown in Figure 1A. We verified that the cost of fluorescence-coding plasmid (i.e. growth rate) was similar for the two sub-strains: The generation times in the exponential growth phase in an M63 minimal medium supplemented with glucose (M63+glu) were (59±1) min and (60±1) min for the GFP and RFP carrying sub-strains, respectively (Supplementary Figure S1).

**Figure 1:**
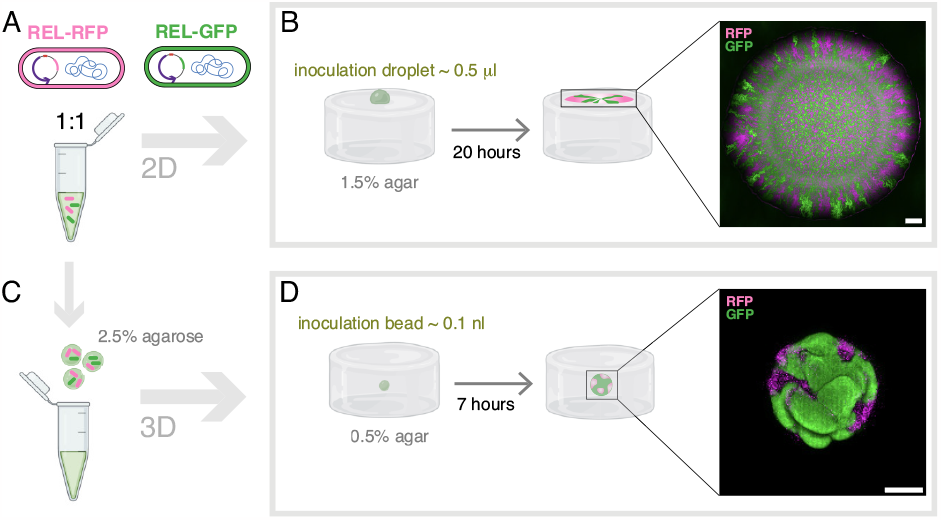
Competition experiments in 2D and 3D. **A:** 1:1 mixture of fluorescent *E*.*coli* REL coding for either GFP or RFP with concentration *c*_0_ or multitudes of *c*_0_. **B:** Sketch of competition experiment in 2D: Inoculation of an 0.5 μl inoculation droplet on M63+glu agar (1.5%) plate, followed by 20 hours of incubation (37°C). The inset is a (maximum intensity) z-projection of an example colony imaged by CLSM. The scale bar corresponds to 50 μm. **C:** Inoculation beads: Small agarose (2.5%) beads encompassing the 1:1 cell mixture from (A). **D:** Sketch of competition experiment in 3D: Inoculation of a 0.1 nl inoculation bead inside a M63+glu agar (0.5%), followed by 7 hours of incubation (37°C). The inset is a (maximum intensity) z-projection of an example colony imaged by CLSM. The scale bar corresponds to 50 μm. Elements of this figure were created with BioRender.com.

When a mixture of bacteria is inoculated on an agar surface, the population expands and segregates into monoclonal sectors. Figure 1B is a sketch of such a (pseudo-)2D competition assay, where 0.5 *μ*l droplets of the 1:1 mix were inoculated on agar (1.5%) supplemented with nutrients. In line with prior findings (Hallatschek et al. 2007), the expansion led to strong de-mixing of the two strains (inset of Figure 1B). In order to set up a 3D version of this experiment, we designed multi-cell beads based on a protocol by Roelof van der Meer and co-workers Buffi et al. 2011. This method, which is sketched in Figure 1C, relies on the hydrophobic nature of silicone oil to form small droplets of an aqueous solution. In our case, this solution is a mixture of bacteria and melted agarose (2.5%). We used this strategy to encapsulate the *E. coli* mixture (Figure 1A) in *inoculation beads*. When embedded in a semi-soft (0.5%) agar supplemented with nutrients and incubated (37°C), colonies outgrow the original agarose sphere to form 3D bacterial colonies as sketched in Figure 1D. The inset is an example of a colony derived from an inoculation bead with 1:1 mixture of fluorescent bacteria of concentration *c*_0_, which corresponds to ∼100 bacteria at the onset of the colony (Table 1).

**Table 1:** Inoculating bead’s concentration in units of OD or cells/ml, using the conversion OD= 1 equal to 1.5 · 10^9^ cells/ml. The 4th column is the calculated number of cells/bead, using the average volume: 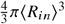 (Figure 2B) and the error correspond to ±SD propagated from the ⟨*R*_*in*_⟩ error.

|  | [OD] | [cells/ml] | [cells/bead] |
| --- | --- | --- | --- |
| $c_0/100$ | $6.9 \cdot 10^{-3}$ | $1.6 \cdot 10^7$ | $1.0 \pm 0.7$ |
| $c_0/10$ | $8.2 \cdot 10^{-2}$ | $1.21 \cdot 10^8$ | $12 \pm 9$ |
| $c_0$ | 0.70 | $1.06 \cdot 10^9$ | $108 \pm 78$ |
| $c_0 \times 10$ | 5.6 | $8.46 \cdot 10^9$ | $863 \pm 625$ |

### Inoculation bead characterisation

Using wide-field fluorescence microscopy, we found that the spherical inoculation beads contain bacteria of both colours, as seen in Figure 2A. We found no bacteria outside the beads, implying that the agarose scaffold protects the cells from the toxic silicone oil (Supplementary Figure S2). Furthermore, we found that their radii, *R*_*in*_, were distributed as given in Figure 2B, with average ⟨*R*_*in*_⟩ = (29±7) μm (mean±SD, *N* = 26), which corresponds to a volume of about 0.1 nl. Hence, the inoculation volume is on the order of 103 times less than in the 2D case (0.5 *μ*l). Our different batches of inoculation beads contained a 1:1 mix of the two colours (GFP and RFP) of bacteria and varying concentrations of cells: *c*_0_× 10, *c*_0_, *c*_0_/10, and *c*_0_/100 (Table 1). In order to estimate the concentration of beads that are able to grow into colonies, we plated the beads on a hard agar surface, incubated the plates, and counted the number of colony-forming units (CFUs). Figure 2C combines the counts fresh from production (opaque) and from frozen stocks (full colour). We found significant differences in CFU per volume for beads of different initial cell concentration. However, in practice we tuned the bead solution to have approximately the same amount of growing colonies pr. culture well.

**Figure 2:**
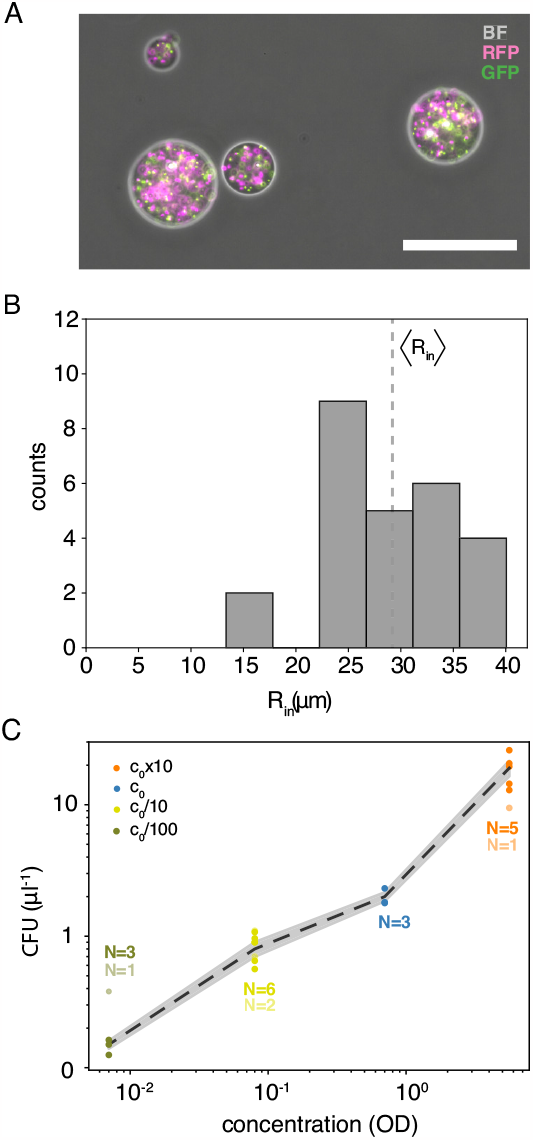
Inoculation bead characterisation. **A:** Inoculation beads with *c*_0_ concentration (Table 1). The image is an overlay of channels: bright-field (BF), green (GFP), and red (RFP). The scale bar is 100 μm. **B:** Distribution of inoculation beads’ radii, *R*_*in*_, with the average value (vertical dotted line):) ⟨*R*_*in*_⟩ = (29±7) μm (mean±SD, *N* = 26) as given in Eq. 1. **C:** Colony-forming units (CFU) corresponding to number of inoculation beads per volume versus density of founder cells, i.e., OD: *c*_0_/100, *c*_0_/10, *c*_0_, and *c*_0_ × 10 on a double-logarithmic scale. Beads are either from frozen stocks (full colour) or fresh from production (opaque), the latter is not included in the mean (punctuated line) and the shaded area corresponds to ±SEM.

### Density of founder cells controls size and patterning

Following agar-embedment and incubation, we imaged colonies using a CLSM. We noted how often we found monocolour colonies (Supplementary Figure S3) but only imaged two-coloured colonies. Figure 3A-C shows three examples of colonies grown from different batches of inoculating beads, i.e., *c*_0/_10 (A), *c*_0_ (B), and *c*_0_×10 (C) and incubated for 7 hours (37°C). At first sight, we found that not only does the size of the colony change with concentration, but also the patterning (more examples in Supplementary Figure S4). For a thorough analysis of this general observation, we segmented and filtered the 3D images to obtain a mask – with the thickness of a single voxel – reconstituting a connected surface and where all voxels were assigned one (and only one) color, see the insets of Figures 4-5 and Supplementary Movie 1 for examples. From this mask we estimated the time-dependent radius, *R*(*t*) for *t* = 7 hours, of the projected surface (assuming spherical symmetry). In line with the general observation, *R*(*t*) grows with the density of founder cells, as shown in Figure 3D.

**Figure 3:**
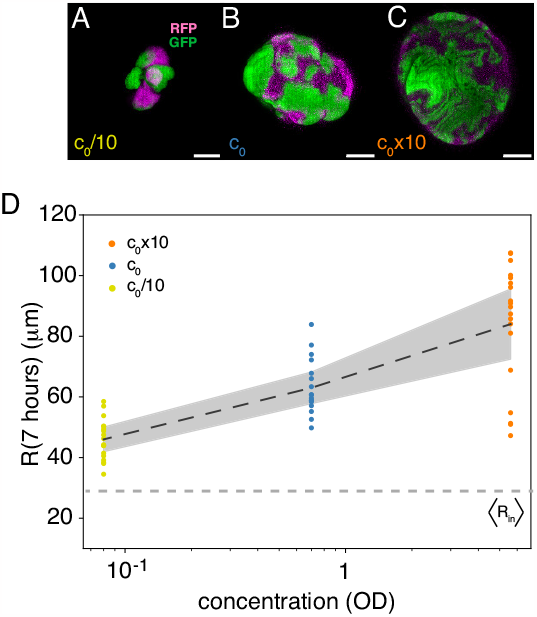
Density of founder cells controls size and patterning. **A-C:** Examples of colonies formed from inoculation beads with different initial concentrations of founder cells: *c*_0_/10 (A), *c*_0_ (B), and *c*_0_ × 10 (C). The scale bars correspond to 50 μm. **D:** Colony radii, *R*(*t*), at *t* = 7 hours for *c*_0_/10 (N=18), *c*_0_ (N=17), and *c*_0_ × 10 (N=19). The punctuated line (dark gray) is the ensemble average (Eq. 1) and the shaded region corresponds to ±SEM. The horizontal punctuated line (light gray) is the average bead size, ⟨*R*_*in*_⟩ (Figure 2B).

**Figure 4:**
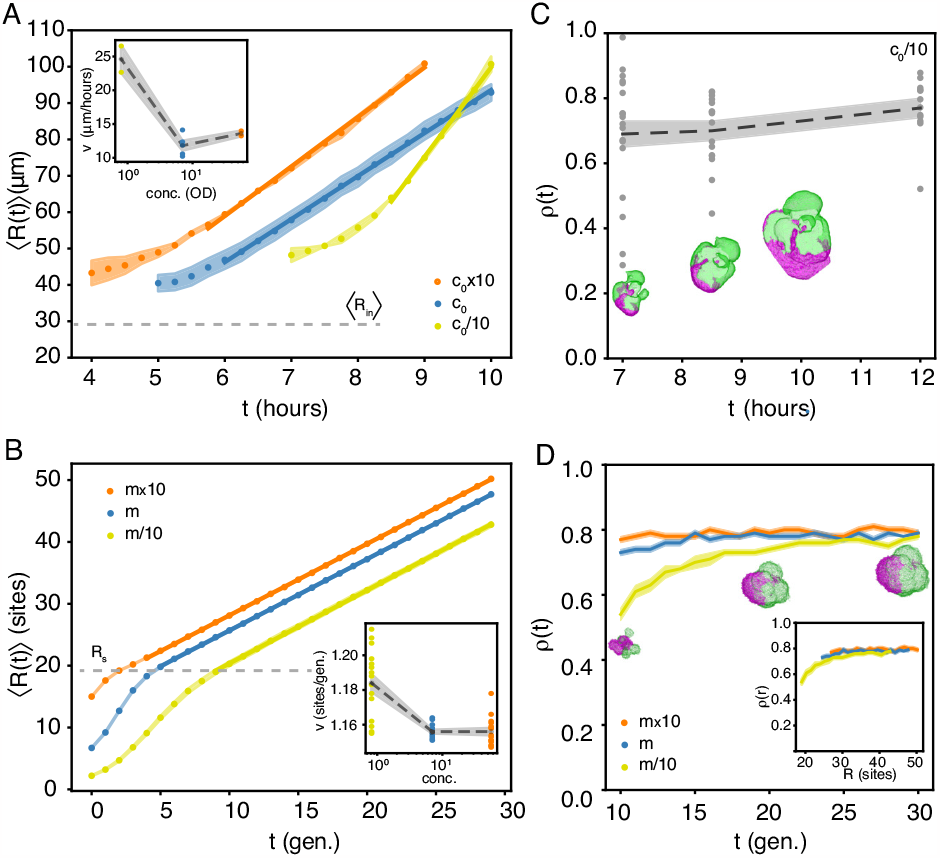
Density of founder cells controls growth dynamics. **A:** Ensemble-averaged colony radii, ⟨ *R* (*t*)⟩, versus time (Eq. 1) for *c*_0_ 10 (*N* = 2), *c*_0_ (*N* = 5), and *c*_0_×10 (*N* = 2). The full line is a linear fit and the shaded area signify ±SEM. The dashed line is the mean radius of the inoculation beads, ⟨ *R*_*in*_ ⟩, found from Figure 2B. Inset: Radial growth speeds, *v*, of the individual traces versus concentration. The dots are the linear fits of the individual *R* (*t*) from the colonies in (A), the dashed are the means, and the shaded area signifies SEM. The dots are the slopes of the individual time-lapses. **B:** Ensemble-averaged *in silico* colony radii, ⟨ *R* (*t*)⟩, versus time (Eq. 1) for *m*/10 (*N* = 15), *m* (*N* = 15), and *m*×10 (*N* = 15). The full line is a linear fit and the shaded area signifies SEM. The dashed line is the mean radius, *R*_*s*_, of the initial seeding sphere (i.e., inoculation beads). Inset: Radial growth speeds, *v*, of the individual traces versus concentration. The punctuated line is the the linear fits of the (full lines) and the shaded area signifies ±SEM. The dots are the slopes of the individual time-lapses. **C:** Isoperimetric quotient, ρ(*t*) (Eq. 3), for different *c*_0_/10 colonies at different times points, *t*: 7 hours (*N* = 18), 8.5 hours (*N* = 16), and 12 hours (*N* = 12). The dashed line is the mean and the shaded area signifies ±SEM. The insets are the masks, *M*, from a time-trace from (A). **D:** Isoperimetric quotient, ρ(*t*), for growing *in silico* colonies (N=15) of concentrations *m*/10, *m*, and *m* × 10 over time, *t*. The full line is the mean and the shaded area signifies ±SEM. The small insets are examples from one model evaluation with concentration *m*/10. Inset: Isoperimetric quotient, ρ(*r*), over radial distance, *R*.

### Density of founder cells controls growth dynamics

To study the dynamics of the 3D bacterial colony formation, we did time-lapses (Supplementary Movie 2) and Figure 4A shows the average radii, *R* ⟨(*t*)⟩, for the three concentrations: *c*_0_ /10 (yellow), *c*_0_ (blue), and *c*_0_×10 (orange) as defined by Eq. 1. The time point at which we began imaging was chosen such that we easily found the small colonies under the microscope (*R* > 40 μm) and we ended at ∼10 hours to prevent out-growing the matrix. Generally, we find an initial growth phase, where the colony radius expands slower than linear over time. This is followed by a temporal window, where ⟨*R*⟩ grows linearly over time (full lines) with the rate, *v*, given in the inset of Figure 4A. This linear expansion indicates that the colonies grow predominantly from the outermost cell layers due to insufficient nutrient penetration deeper in the colony Kuennen and Wang 2008. Also, this is well in accordance with earlier predictions Shao et al. 2017, Lavrentovich and Nelson 2015 and experimental findings for a similar *E. coli* system Cordero et al. 2023. Notably, from these earlier predictions, we would expect *v* to be similar for all concentrations, as is the case for *c*_0_ and *c*_0_ × 10. However, our results from the lowest density of founder cells (*c*_0_/10) contradict this view (inset of Figure 4A). Instead, the radial range expansion rate is significantly faster (*v* ∼ 25 μm/hour vs. ∼ 13 μm/hour). Assuming the doubling time to be ∼ 1 hour (also inside 0.5% agar) and the volume of a single cell to be ∼ 1 μm^3^, this corresponds to a growing layer of the thickness of about 25 cells.

As it is unclear how fewer inoculating cells result in faster radial growth, we explored this question employing an Eden growth lattice model initiated from two species of seeds/cells with identical properties randomly placed in a sphere of radius, *R*_*s*_, of three different concentrations: *m*/10, *m*, and *m*×10 (Materials and Methods). These seeds were allowed to divide into empty unoccupied neighbouring sites until they eventually grew as one cluster Jullien and Botet 1985. The resulting pattern mimicked the competition of two populations in confined bacterial colonies. To study the dynamics of the resulting *in silico* colony formation (Supplementary Figure S5), we created time-lapse movies of growing colonies with different seeding concentrations (Supplementary Movie 3-5). Figure 4B shows the average radii, ⟨ *R* (*t*)⟩, for the three concentrations: *m*/10 (yellow), *m* (blue), and *m*×10 (orange) in the linear growth part of the curve as defined by Eq. 1. In contrast to the experimental observation, we observed the initial growth (⟨*R*(*t*)⟩ < *R*_*s*_) in the simulation to be faster and settling to linear growth with a constant rate (inset of Figure 4B). We find large variations in *v*, especially for the lowest seeding concentration (*m*/10). Moreover, the radial velocity is largest for this concentration (even though the difference is small) because of rougher surfaces.

In the Eden growth model, the expansion speed is dominated by the number of cells on the surface (i.e., cells neighbouring empty lattice sites). Because colony shapes are rougher for lower concentrations of founder cells and for earlier times (Figure 4D), the radial growth at earlier times will be faster. This effect is especially visible when the colony is smaller than the seeding sphere (⟨*R* (*t*)⟩ ≤ *R*_*s*_) and the occupied lattice sites are not yet fully connected. Moreover, the colony’s morphology could also cause fast growth of the colonies initiated from *c*_0_/10 inoculation beads in the experiment. In particular, protrusions on the surface enlarge the surface-to-volume ratio and may enhance growth by making nutrients accessible to a larger number of cells. To investigate this, we measured the time-dependent isoperimetric quotient, ρ(*t*) (Eq. 3), for the projection of *c*_0_/10 colonies, as shown in Figure 4C (Materials and Methods). ρ(*t*) is a dimensionless measure of compactness and approaches its maximum for a perfect circle projection. We found large scattering in ρ(*t*), especially for small *t*, but the average ρ(*t*) tends to grow slightly, as the cracks and valleys are filled and colonies become more round. For comparison, in Figure 4D, we measured ρ(*t*) for the modelled data for *t* ≥ 10 generations, when the colony is one connected structure and ⟨*R*(*t*)⟩ shows linear behaviour over time (Figure 4B). For the modelled data, we found that ρ(*R*) is independent of inoculation density (inset Figure 4D) and that it converges to a value comparable to the experimentally observed ρ(*t*).

### Occupancy on colony surfaces matches the initial ratio in the inoculation bead

In order to evaluate any competitive advantage for one sub-population over the other, we measured the occupancy, 𝒪 7 hours, or the fraction of the GFP-expressing sub-strain on the colony surface (Eq. 4). We found equal occupancy of two sub-populations, as the fraction of the surface occupied by GFP-carrying cells is centred around 𝒪 = 1/2 for all inoculation bead concentrations (Supplementary Figure S6). As the sub-populations have equal fitness advantage (Supplementary Figure S1), the final surface ratio corresponds to the 1:1 mixture in the inoculation bead. Also, we find from our model that time-dependent fluctuations in 𝒪 (*t*) are more important for smaller inoculation concentrations (Supplementary Figure S7).

### Sector patterns are dynamic

In aim of characterising the sectors on the colony surfaces and their time-dependence, we identified the different sectors on the 3D *in silico* colony surfaces, as green and magenta in the examples in Figure 5A. Then we calculated their time-dependent areas, *A*_*sec*_ (*t*), on the surface (Materials and Methods). From the cumulative frequency, as shown Figure 5B, at a specific time point (*t* = 29 gen.), we found sectors to be larger for low seeding density. With this version of the Eden growth model, we loose track of the individual lineages. Therefore, we ran a modified model in parallel (Supplementary Figure S8 and Supplementary Movie 6-8), which grew from multi-seeds (Materials and Methods). From the cumulative frequencies (inset Figure 5B) we found that the majority of sectors are smaller than 10 sites and that the largest sectors indeed are merges of sectors of different lineages. We complemented this finding with the *A*_*sec*_ (*t*) at *t* = 7 hours for both *c*_0_ /10 and *c*_0_-derived colonies, as shown in Figure 5C. Even though our detection limit was 10 voxels (∼370 μm^2^), we found that smaller sectors dominated. We left *c*_0_×10 colonies out of the analysis, since many sectors were smaller than our detection limit and resolving colours was difficult (Supplementary Figure S9).

**Figure 5:**
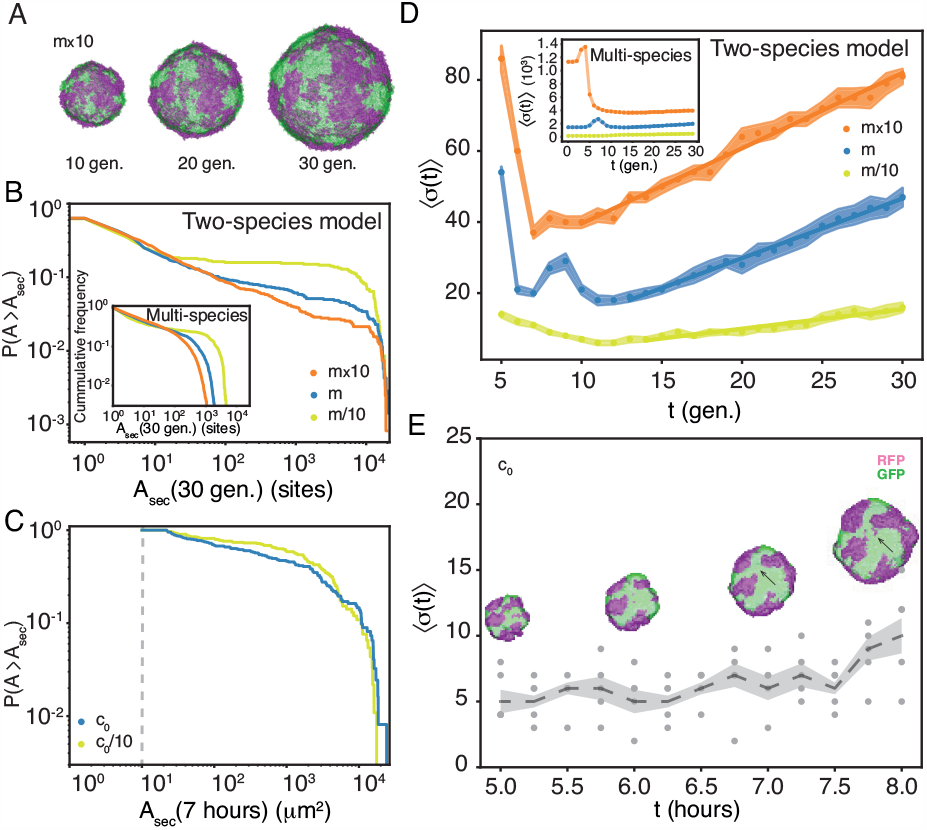
Sector patterns are dynamic. **A:** An example of how surface sectors changes with time, *t* ∈ {10, 20, 30} generations in a two-species *m*×10-derived modelled colony. **B:** Cumulative distributions of areas of sectors,, *P*(*A*≥ *A*_*s*_*ec*), of sub-population on the two-species *in silico* colonies’ surfaces, *A*_*sec*_ (*t*), at *t* = 30 generations for *m* /10 (*N* = 233), *m* (*N* = 702), *m* ×10 (*N* = 1219) sector from 15 different colonies. Inset: Multi-species model for *t* = 29 gen. *m* 10 (*N* = 708), *m* (*N* = 2978), *m*×10 (*N* = 5939) sectors from 15 different colonies. **C:** Cumulative distributions of areas of sectors, *P* (*A*≥ *A*_*s*_*ec*), of sub-population on the experimental colonies’ surfaces, *A*_*sec*_ (*t*), at *t* = 7 hours for *c*_0_/10 (*N* = 92) sectors, as well as *c*_0_ (*N* = 123) sectors from 18 and 17 colonies, respectively. The vertical dashed line signifies the detection limit of 10 voxels. **D:** Ensemble-averaged number of sectors, ⟨σ (*t*)⟩, versus time, *t*, (Eq. 5) for concentration: *m* /10 (yellow), *m* (blue), and *m* × 10 (orange) as given in Eq. 5. There is no distinction between the two-colors of sectors and the number of colonies is (*N* = 15). The splitting rate found by linear fit of the linear part is 0.55 ±0.04 gen^−1^, 1.68 ±0.04 gen^−1^, and 2.12 ±0.06 gen^−1^ for *m*/10, *m, m* × 10, respectively. Meanwhile, the density of sectors reaches a constant low level (Supplementary Figure S8A). The shaded area signifies SEM. Inset: The same for the multi-species model, where the splitting rate is 1.03 ± 0.07 gen^−1^, 3.56 ±0.13 gen^−1^, and 2.35 ±0.17 gen^−1^ for *m*/10, *m, m* × 10, respectively. Also for this model, the density of sectors reaches a constant low level (Supplementary Figure S8B). **E:** Ensemble-averaged number of sectors, ⟨σ (*t*)⟩, versus time, *t*, (dashed line) as given in Eq. 5 for *c*_0_ (*N* = 5). The dots are the number of sectors for individual colonies without distinguishing between red and green. The shaded area signifies ±SEM. Inset: The time evolution of surface sectors for a *c*_0_-derived example colony. An example of sector splitting is designated (black arrows).

We sat out to investigate the time-dependence of the number of sectors, *σ* (*t*), using our Eden growth models. As shown in Figure 5D, the average ⟨*σ*(*t*)⟩ first decreases, as a consequence of merging of clusters from different founder cells (note that we counted spatially separated clusters as different sectors), followed by a linear increase in sectors, due to sector splitting. The merging (i.e., ⟨σ (*t*)⟩ decrease) is even more pronounced for the multi-species model (inset Figure 5), even though sub-sequent splitting rates are larger.

We finally looked into how the number of sectors changes with time, *σ*(*t*), in *c*_0_-derived colonies in Figure 5C. Even though we expected more splitting events for *c*_0_ × 10, again, the most sectors were below our resolution. Based on the time-lapses of 5 colonies, we found that the ensemble-average, ⟨*σ*(*t*)⟩, as defined in Eq. 5, goes slightly up within our time range. This corresponds to a splitting of sectors, as shown for one example in the inset of 5E (black arrow). In contrast, the sector merging was not possible to observe experimentally since it happened inside the beads right after embedment (*t* < 5 hours) and we were limited by our optical resolution. Once the clusters are connected, the cells grow as one colony and hereafter sector splitting dominates giving an increase in⟨σ (*t*)⟩. It is worth noticing that there is no obvious way to calibrate the experimental timescale against the simulation timescale, as we ignore the spatial interpretation of the model’s lattice sites.

## DISCUSSION

Recent advances in 3D culture technology allow for *in vitro* models of development and disease with organoids (i.e., simplified organs) Bredenoord et al. 2017 or tumorospheres Weiswald et al. 2015, Costa et al. 2016, respectively. The counterpart of these 3D models within bacteriology research is quasi-spherical colonies, grown from a few bacteria embedded in agar. We propose 3D cultures initiated from agarose droplets could be this simple (yet highly controllable) multi-species model of biofilm-formation in soil, natural water environments, minerals, or other substrates.

In the present manuscript, we presented a method, which in brief, substitutes the inoculation droplet with an inoculation bead of much smaller volume (fractions of *μ*l versus nl). So, our colonies are initiated from very small volumes and we can, hence, inquire about the early stages of colony development. Furthermore, we interchanged the hard agar (> 1%) substrate with submersion in soft agar (< 1%), which allows for studies of spherical range expansion. We used confocal laser-scanning microscopy (CLSM) for imaging but other techniques to optimise the resolution and light penetration in large cell agglomerates are evolving rapidly (e.g light-sheet microscopy Tomer et al. 2012). Even though advanced imaging is required for proper 3D reconstitution, we anticipate that many questions could be answered by 3D competition assays under wide-field fluorescence microscopes.

From 2D competition experiments, we know that sector boundaries are diffusive Hallatschek et al. 2007 and for rod-shaped cells even super-diffusive Hallatschek and Nelson 2010. The reason for this is that uni-axial growth of rod-shaped bacteria (i.e. *E. coli*) results in chain-formation of bacteria. As compressing forces make small asymmetries in cell alignment, these instabilities propagate to cause jagged shapes on surface-attached monolayers Boyer et al. 2011, Rudge et al. 2013. These shapes are further enhanced by inter-cellular adhesion, which leads to increased diffusivity of mixing and area of interaction between lineages Kan et al. 2018. We find similar patterns of jagged sector boundaries on the colonies’ surfaces (Figure 3C), even though our boundaries are 2D interfaces between sectors. This highlights an interesting following-up question: how does cell alignment affect the spatio-temporal dynamics of sectors?

The Eden growth model predicts a transition from super-linear growth in early colonies to slower (i.e. linear) surface expansion over time (Figure 4B). The super-linear regime largely overlaps with the regime where the growth – starting from the individual founder cells – is not yet connected into one colony. Our experimental resolution did not allow us to observe such early super-linear dynamics and instead, we observed a slower growth. We speculate that this slow growth could be due to i) a possible heat shock from taking the culture wells out from the incubator to the CLSM for imaging the colonies and/or the higher mechanical stress the cells might be under when growing insie the agar beads (2.5% agarose) and that this stress reduces once cells have outgrown the beads and expand in the agar matrix, allowing them to growth faster. We did observe the linear surface expansion in experiments. The reason for this linear regime is that behind the fast-growing surface cells, is a quiescent region, where proliferation is slower because of space constraints Tuson et al. 2012 and metabolite diffusion from the colony periphery Shao et al. 2017, Lavrentovich and Nelson 2015, Lavrentovich et al. 2013a. We also find that the entrance into the linear regime depends on the initial concentration of founder cells, where the colonies with the highest initial concentration reach this compact-mature colony state earlier (Figure 4A). This is also consistent with the simulations (Figure 4B).

We also found that, for the lowest density of founder cells, the colonies have a rougher surface growth as earlier reported Martínez-Calvo et al. 2022. The cells located in these bumps or protuberances are expected to have better availability of nutrients, which can be part of the explanation for their observed faster radial growth (inset Figure 4A). The model also showed slightly faster growth for lower initial cell density (inset Figure 4B), though the difference was small. Furthermore, we believe that the surface roughness decreases slowly with time and the colonies become rounder and rounder (Figure 4C). For 2D colony growth, it has been established that colony morphology qualitatively changes with nutrient availability and agar hardness. Specifically, nutrient-poor environment and/or harder agar can cause instabilities to develop branching Ohgiwari et al. 1992. Similar instabilities are expected in 3D growth under certain environmental conditions. The observed lack of strong instabilities in our system supports the use of the Eden growth model that ignores inhomogeneity and nutrient depletion of the environment. Our model presumes homogeneity inside the colony. We speculate that the merging of somewhat grown microcolonies may lead to structural variations and affect the overall growth. Hence, to understand the growth dynamics quantitatively, a model that considers these factors would be desirable.

We have demonstrated that the sectors on the surface can merge but also split over time. In our experiments, we observed the splitting events, and the number of sectors showed a tendency to increase, though fluctuation was large (Figure 5C). These observations are consistent with the multi-species Eden growth model simulation and is independent on motility of individual cell (as our strain lacks flagella Studier et al. 2009). In 3D expanding colonies, individual clusters – starting from individual seeds – quickly merge to form a connected colony, and after that, although large sectors do form, they also fragment forming small sectors. Therefore, the number of sectors actually increases over time (Figure 5E). This contrasts what we know from 2D radial expansions, where an initial coarsening phase, in which domain boundaries annihilate, is followed by a stable phase, in which the number of sectors does not change Hallatschek et al. 2007. This behaviour in the 2D system is the consequence of initial merging of sectors, due to the diffusion of sector boundaries and the expansion of the radial frontier, which is inflating the distance between the sector boundaries. This results in the crossover time *t*^*\**^ ∼*R*_0_/*v* after which the coarsening is suppressed Lavrentovich et al. 2013b. Here, *R*_0_ is the radius of the initial “homeland” which determines the length scale of the initial sectors, and *v* is the expansion speed of the radius. In 3D colony growth, the sector boundary on the 2D surface is a fluctuating line, and a new domain can “pinch off” to form a new sector (e.g. Castillo and Lavrentovich 2020). When we ignore the effect of the roughness of the surface, the surface dynamics of the two-species Eden growth model are expected to show the same dynamics as the voter model Lavrentovich and Nelson 2015, Lavrentovich et al. 2013b. For the voter model on a flat surface, the coarsening dynamics over time, *t*, is marginal, where the density of the boundary between the sectors decreases in proportion to 1/ln *t*. This is because there is no “surface tension” to keep the sector boundaries compact. Even though merging of sectors happen, the boundary of a large sector can fluctuate locally and segment into small sectors Dornic et al. 2001. In the spherical expansion, the increase of the surface area is expected to suppress the already marginal coarsening. It should be noted, however, that the total surface area is increasing proportional to *t*^2^. The number of sectors per area (i.e., surface density) in the Eden growth model showed a decreasing tendency over *t* especially for the high-seed concentration (Supplementary Figure S10). This suggests that the expansion of existing sectors dominates most of the surface. In addition, it is worth noting that the roughness of the surface can affect the coarsening dynamics Kuhr et al. 2011. However, further theoretical studies are needed to characterise mixing and de-mixing dynamics in 3D range expansion quantitatively.

As our method is both cheap and easy to set up, this can be a natural extension of the original 2D competition assay to study 3D dynamics. The method can also provide a platform for a broad range of investigations with the potential for high throughput experiments Friedrich et al. 2009. Furthermore, this model system could be used to resolve, not only how bacterial competition drives spatio-genetic patterning in neutral competition, but also how selective advantage, competitive interaction, cooperation, and division-of-labor among the different cells distort this pattern in 3D growth by introducing different interacting cell types Lee et al. 2022, Dal Co et al. 2020, Borer et al. 2020, Celik Ozgen et al. 2018. Therefore, we anticipate that the proposed accessible method can help advance our understanding of microbial community development and evolution in related systems in 3D environments significantly.

## MATERIALS AND METHODS

### Bacterial strains and culture media

We used two sub-populations of the *Escherichia coli* B strain REL606 Monds et al. 2014 carrying plasmids that constitutively expressed kanamycin resistence and either green fluorescent protein (GFP), pmaxGFP (pmaxCloning-Vector, Lonza), with excitation/emission 487/509 nm or red fluorescent protein (RFP), pTurboRFP (pmaxCloning-Vector, Lonza), with 553/574 nm Smith et al. 2017. Therefore all media used in this study was supplemented with 30 μg/ml kanamycin to retain fluorescence.

Through this study we alternated between the following two medias: the rich Lysogenic broth (LB) with 1% Bactotryptone, 0.5% NaCl, and 0.5% yeast extract and the M63 minimal media (M63+glu) with 20% 5×M63 salt Elbing and Brent 2002, 1 μg/ml B1, 2 mM MgSO_4_ and 2 mg/ml (w/v) glucose.

### Encapsulating bacteria in inoculation beads

For a 3D competition experiment, we produced 2.5% agarose beads following the procedure of Ref. Buffi et al. 2011. One day prior to bead-formation, pipette tips were placed in an incubator set to 60°C, two overnight cultures of REL+GFP and REL+RFP in 2 ml of LB were prepared, and incubated at 37°C under constant shaking. The following day, we measured (NanoPhotometer C40) the optical densities at the wavelength of 600 nm (OD) of the overnight cultures. Then, we prepared a saturated 1:1 mixture, by mixing the overnight cultures in the ratio corresponding to their ratio of the measured ODs. Meanwhile, 15 ml of silicone oil (dime-thylpolysiloxane, Sigma) in a 50 ml tube was heated in a block heater at ∼55°C (AccuBlock Digital Dry Baths, Labnet International, Inc.) for ∼20 min. Then, sterile 2.5% agarose (VWR Life Science, EC no:232-731-8, CAS No:9012-36-6) in Mili-Q water was melted in a microwave. We made sure to i) shake rigorously for homogeneity and ii) avoid boiling to minimise evaporation. Then, the agarose solution was left to cool to 55°C, before 500 *μ*l was transferred to a 2 ml (heated) tube in the block heater (55°C). Afterwards, 15 *μ*l of pluronic acid (Pluronic F-68 solution 10% Sigma) was added and the tube was transferred to a shaker block heater (Thermometer Confort, Eppendorf) set at 42°C. After few minutes, 2-100 *μ*l of the saturated 1:1 bacteria culture was added under continuous vortexing (1400 rpm) for final concentrations of OD ∈ {5.6, 0.70, 8.2 ·10^−2^, 6.9 ·10^−3^} (Table 1). We removed the silicone oil from the block heater and – using the pre-heated pipette tips – 500 *μ*l of this bacteria-agarose-pluronic acid mix was added drop by drop to the silicone oil. This step was done fast to avoid untimely solidification of the agarose (at temperatures < 25°C). Afterwards, the 50 ml tube containing the droplets in silicone oil was vortexed (Vortex Mixer, Labnet International) for 2 min at maximum speed before being placed in a water-ice bath for additional 10 min. Then, centrifuged for 10 min at 550 g (room temperature), before the silicone oil was removed by careful pipetting followed by 2×washing: i) Addition of 5 ml of phosphate buffered saline (PBS), ii) centrifugation for 10 min at 550 g, and iii) gentle removal of oil and supernatant. Finally, the washed beads were resuspended (by vortexing) in 5 ml of PBS.

To harvest the beads, we let bead-PBS solution flow through a set of strainers with mesh sizes of 70 μm (Cell strainer 70 μm Nylon, BD Falcon) and 40 μm (Cell strainer 40 μm Nylon, BD Falcon), placed on a 50 ml tube. An additional 5 ml of PBS was poured through the strainers to ease the passage. To collect the selected beads, the 40 μm strainer was placed upside down on a new 50 ml sterile tube and the beads adsorped to the filter surface were desorbed by rinsing with 6 ml of LB medium. For long-term storage, the bead-medium solution was aliquoted in volumes of 500 *μ*l in 2 ml vials, before mixing with 500 *μ*l of 50% glycerol-solution (11 vials pr. batch), and stored at -80°C. We used 1 ml serological pipette tips for bead handling to prevent damaging flow rates.

### Growth rate measurements

Cultures of REL+GFP and REL+RFP were incubated (37°C) in parallel overnight in 2 ml of M63+glu under constant shaking. The following day, 500 *μ*l of each overnight culture were diluted in 5 ml of M63+glu and the ODs were measured over 5 hours with a sampling rate of ∼ 30 minutes.

### Inoculation bead concentration measurements

#### Ratio of coloured strains

To verify that the ratio in the beads (tuned with OD ratios), we serially diluted (10^6^ ×) the saturated 1:1 cell mixture (for encapsulation) and plated (50-200 *μ*l) on LB plates with agar (1.5%). The plates were incubated overnight, and the number of CFUs from REL+RFP and REL+GFP was counted to verify the 1:1 ratio (Supplementary Figure S11).

#### Cells pr. bead

Overnight cultures of REL+GFP and REL+RFP was serially diluted (10^6^ ×) and plated (50 *μ*l) on LB plates with 1.5% agar. From the number of CFU, we found that OD= 1 corresponds to ∼1.5 ·10^9^ cells/ml. Combined with OD∈ { 5.6, 0.70, 8.2·10^−2^, 6.9·10^−3^} and the average bead volume, we estimated the average number of cells pr. bead given in Table 1.

#### Beads pr. volume

We mixed 80 *μ*l of inoculation beads, either directly from production or from frozen vials, with 200 *μ*l medium (LB). The mixture was plated by gently whirling (no spatula) on LB plates with 1.5% agar, then the plates were incubated overnight, and finally the number of CFUs was counted. This estimation of beads pr. volume was used to determine the proper dilution after thawing to ultimately get ∼ 20 beads pr. well.

### Imaging inoculation beads

To estimate inoculation beads’ radii, beads were imaged with an inverted Nikon Eclipse Ti fluorescent microscope (Nikon, Tokyo, Japan) using a 20× air immersion objective (Splan flour,L20×,0.45corr ∞) paired with an Andor Neo camera (Andor, Belfast, UK). GFP and RFP was excited by a Hg lamp using the FITC and Texas-red (Nikon, Tokyo, Japan) cubes, respectively.

### Toxicity measurements

To evaluate the toxicity of silicone oil and pluronic acid, we did limiting dilution experiments: Overnight cultures of REL+GFP and REL+RFP was (10 - 10^7^)× diluted in 1 ml of either pluronic acid, silicone oil, or PBS as control. Then the number of viable cells was determined by plating (LB plates with 1.5% agar) followed by CFU counting.

### Competition assays

#### Embedment of inoculation beads in 0.5% agar

In line with methods detailed in Cordero et al. 2023 and references therein, bottles of 20 ml milipore water with 0.625% agar were melted by repeated cycles of heating (in the microwave) and shaking to ensure homogeneity and minimise evaporation and ageing Mao et al. 2017. After cooling (∼55°C), we added M63 salt Elbing and Brent 2002 and supplements to a final M63+glu with 0.5% agar. Then, 1 ml of this solution was transferred to a pre-heated tube in the (non-shaking) block heater (55°C). A frozen vial of bead-medium solution was thawed (on the bench) for ∼5 min until melted and 20-500 *μ*l were transferred into 1 ml of LB (to minimise the time spend in high-concentration glycerol). Meanwhile, the agar-medium solution was supplemented with 10 *μ*l of a 1 M stock of KNO_3_ – to avoid growth difficulties in anoxic conditions as suggested by Refs. Sønderholm et al. 2017, Kvich et al. 2019 – and 30 μg/ml kanamycin. Quickly hereafter, 50 *μ*l of the diluted bead-medium solution was added and the tube’s content was mixed and poured into a glass bottomed culture well (WillCo HBST-5040). Here it was left to solidify for a few minutes on the bench before incubation (upside-down) at 37°C for at least 5 hours (see imaging section). The agar had a final thickness of ∼300 μm and contained 17±5 inoculation beads/well (Supplementary Figure S12); corresponding to at the most one bead for every 50 *μ*l agar. For 3D competition experiments, the wells were incubated at 37°C for 5-7 hours (depending on the experiment).

#### Deposition of inoculation droplets on 1.5% agar

In line with the protocol of Ref. Jauffred et al. 2017 and references therein, we mixed 1:1 overnight cultures of REL+GFP and REL+RFP with LB to a final OD=0.7 before inoculating 0.5 *μ*l on plates with M63+glu and 1.5% agar. Plates were incubated for 20 hours at 37°C.

### Imaging competition assays

#### 3D colony images

After incubation, we checked all culture wells to discard those, where colonies had outgrown the agar and spread over the agar-air interface. All colonies of the remaining culture wells were imaged with CLSM (Leica SP5) and a 20× air objective (NplanL20×0.40NA). The data was acquired with sequential z-stacks of RFP (488 nm laser, detection range of (498-536) nm) followed by GFP (543 laser, detection range of (568-641) nm) to a final image of voxel size in (x,y,z) of (1.51,1.51,1.33) μm. For every new condition, we made sure to collect images from at least 2-3 different wells.

#### Time-lapse 3D images

After 4-7 hours of incubation, our colonies had outgrown the inoculation beads and reached a diameter of ∼100 μm. For CLSM time-lapses, our microscope was equipped with a standard water-based heating stage set to 37°C and we did 5 hours of imaging with a frame rate of 4/hour. We repeated this 2-5 times for each condition. We also investigated time-evolvement by comparing individual sets of colonies incubated for different amount of times. The visualisation was made using the Fiji 3D viewer plugin Schmid et al. 2010.

#### 2D colony images

The agar plate was placed up-side down on our inverted CLSM. They were imaged with sequential z-stacks using a similar procedure as for the 3D colonies but with an 5× air objective (Nplan5×0.12PHO, Leica). The voxel size in (x,y,z) was (6.07,6.07,10.33) μm and we used Pairwise Stitching Preibisch et al. 2009 plugin for Fiji.

### Image analysis

#### Segmentation of 3D colonies

The 3D image segmentation was done using both BiofilmQ Hartmann et al. 2021, Matlab, and Fiji/Image J Schindelin et al. 2012 and the following work flow. First, the segmentation was done separately for each channel (GFP+RFP) in BiofilmQ. We cropped the image stacks in z-direction (same *z* for both channels) to discard the colony half-sphere furthest away from the objective, which was distorted by strong shadowing effects. Then, we de-noised the images by convolution ([kernel size: [x,y,z] = [5,5,3] pixels) to soften the edges and fill the holes between cells. Using the Otsu method (with 2 classes: object and background), we manually selected the threshold value for each image in the stacks. Afterwards, we checked – using the Overlay function – that the segmentation matched the raw data. Hereafter, the resulting masks of the channels (GFP+RFP) were divided into small cubic volumes of length 1.52 μm, which reflects the voxel size of the image and corresponds 2-4 *E. coli* volumes. Hereafter, the channel masks were merged, and the surface of the resulting mask was found calling Cube_Surface followed by the Filter Objects function with Cube_Surface in the range [1,2]. The resulting segmentation of the merged channels were saved (mat-file) and the Cube_Surface voxel indices vector was converted to a matrix (using Cell2mat). The matrix can be read as a 3D logical image of a connected surface, but what we wanted was a hollow half-sphere with the thickness of a single voxel. Therefore, we manually removed the surface furthest away from the objective using the Paintbrush tool in Fiji.

The resulting segmentation mask, was multiplied separately with each segmented channels (GFP and RFP) and then merged to the resulting surface mask. We found the number of voxels in the surface mask that was assigned both colours (GFP and RFP), to be (3±2) %, (4±1) % and (11± 4) % for *c*_0_ /10, *c*_0_, and *c*_0_×10, respectively (Supplementary Figure S9). As they are few and that RFP in our system is less detectable, we decided to assign all these two-coloured voxels to RFP. We also found that the surface mask had some discontinuities (<), using 3D Fill Holes plugin in Fiji Ollion et al. 2013. Hence, the resulting surface mask, *M*, is a single layer of voxels each with one and only one colour (see an example in the Supplementary Movie 1).

#### Size estimation of 3D colonies

From z-projections of the mask *M* (or the thresholded image of beads), we obtained logical (x,y)-images. The projected areas of the *n*-th colony (bead), 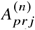, was found by counting pixels ≠ 0 and multiplying by pixel area conversion in (x,y). By assuming the colony (bead) to be spherical, we estimated the ensemble-averaged radius, *R*, at time *t* to be:

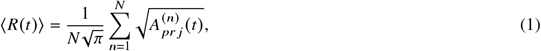

where *N* is the total number of colonies.

#### Determining perimeter in 3D bacterial colonies

From z-projections of the mask *M* (or the manual thresholded images of the z-projections), the projected perimeter, *P*, of the *n*-th colony was found by using the Fiji function called Analyze Particles.

#### Isoperimetric quotient

Measuring the perimeter, *P*, of *A*_*pr j*_ (*t*) for a projection of a colony we found the ratio between *A*_*pr j*_ (*t*) and the area of a circle with the same *P*, called the isoperimetric quotient for a colony to be:

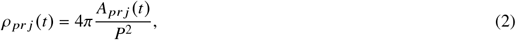

which is a dimensionless measure of compactness with the maximum compactness for a perfect circle (ρ = 1). However, because of square pixels that roughens the shape boundaries, we normalised each value of ρ _*pr j*_ (*t*) with the ρ (*t*)^′^ corresponding to a pixel-resolved circle with an area ∼*A*_*pr j*_ (*t*). In detail, we found ρ (*t*) ^′^ of a circle with pixelated boundaries circle with the area: *A*^′^ = *A*_*pr j*_ and perimeter *P*^′^. The result is then

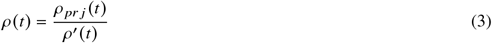

#### Determining occupancy in 3D bacterial colonies

We defined the ensemble-averaged occupancy of GFP-expressing bacteria, 𝒪, on the colony surface as the sum over the following ratio:

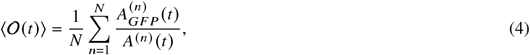

where 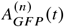 is the area of GFP-expressing bacteria and *A*^(*n*)^ (*t*) is the total surface area of the *n*-th colony at time *t*, such that 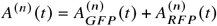.

#### Sectors in 3D bacterial colonies

We defined 3D sectors as connected regions on the colony surface using the Fiji plugin Find Connected Regions on the colony mask *M* with a minimum sector area of 10 voxels (∼23 μm^2^). This value was chosen to be optimal for our dataset by comparing the automated sector counts, *σ*(*t*), with manual counts. The area of a sector (GFP or RFP) at a given *t* is *A*_*sec*_ (*t*), which we found by scaling the number of voxels in the sector with the conversion in (x,y). We plotted them as so-called survival curves, which is the cumulative probability of finding *A*_*sec*_ (*t*) (for a specific *t*) larger than a specific value of the area, *a*: 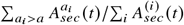, where *i* is the integer of all sectors in all colonies.

We also found the ensemble-averaged number of connected sectors (GFP+RFP) on the colony surface:

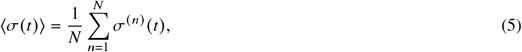

where *σ*^(*n*)^ (*t*) is the number of sectors on the colony mask of the *n*-th colony at a given *t*.

For the modelled colonies we used the minimum sector area of 1 lattice site, as there is no ambiguity in the assignment of colours in this case.

### 3D Eden growth model

We employed 3D Eden growth model to simulate bacterial colonies grow from an inoculation bead in a 3D environment. The model divides the space into a cubic lattice, where each lattice site can be occupied by “bacteria” or empty, and the growth happens if there is an empty neighbor site. It should be noted that the correspondence of the length and time scale between the model and the experiment is only qualitative; this is a coarse-grained model where one occupied lattice site represents a meta-population of a cluster of several cells Fusco et al. 2016.

#### Initial condition two-species population

In a 3D cubic-lattice of size (*L* × *L* × *L*) with *L* = 200, we placed (in the centre) a sphere of radius, *R*_*s*_ = 19 lattice sites. Individual seeds were placed randomly inside the sphere and the number of seeds were on average N∈ { 16, 144, 1150} drawn from a 1:1 distribution of the two colours of seeds (1 or 2), corresponding to seeding concentrations of *m*/10, *m*, and *m*×10. More specifically, we filled each site inside the sphere with the probability *N*/*V* with 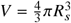, with one of the two colours assigned with equal probability.

#### Initial condition multi-species population

The initial conditions were the same as for the two population model, except that each seed was different: [1,., ⟨*N*⟩] with equal probability (*N*/*V*).

#### Update rule

We first make a list *S* containing all surface sites, i.e., occupied sites with one or more empty neighbouring sites out of the 6 neighbours. The growth is done as follows: (i) Choose one site to divide from the sites in the list *S* randomly with equal probability. (ii) Randomly choose one of the empty neighbour sites of the chosen site in (i). (iii) Fill the chosen site in (ii) with the same color as the chosen site in (i). (iv) Update the list *S* such that a) the new site was added to *S* if it had one or more empty neighbouring sites and b) the sites were removed from *S* if they no longer had any empty neighbouring sites. We repeated (i)-(iv) 5·10^5^ times. For every division, the time proceeds by the inverse of the number of surface sites in *S*, meaning that a unit of time corresponds to the “generation time” (an occupied site on the surface duplicates on average once per unit of time).

We coded the model in C++ using a C++ library Armadillo (Sanderson and Curtin 2016).

### Statistics

All mean values are given as mean plus/minus the standard error of the mean (SEM) or the standard deviation (SD) and only when data are tested against the null hypothesis that it is normally distributed. For distributions, bin sizes were chosen following Sturge’s rule, unless stated otherwise.

## ACKNOWLEDGEMENTS AND FUNDING SOURCES

The authors thank Martin Møller Larsen for fruitful discussions on the modelling and for providing parts of the code for the modelling. The authors also thank Thu Trang Nguyen for providing the conversion factor from optical density to number of cells (colony forming units) and Nathánaël van den Berg for assistance in the competition experiment in 2D.

This work was funded by Danmarks Frie Forskningsfond grant DFF0165-00032B and DFF0165-00103B (LJ) and by Novo Nordisk Fonden grant NNF21OC0068775 (NM).

## DATA AVAILABILITY STATEMENT

All data and detailed protocols are available upon reasonable request and code is available on Zenodo (doi:10.5281/ZENODO.8245914).

## Notes

https://zenodo.org/records/8245914

### Competing Interest Statement

The authors have declared no competing interest.

### Summary of Updates

To accord for typos and small clarifications

## REFERENCES

Eigentler, L., F. A. Davidson, and N. R. Stanley-Wall, 2022. Mechanisms driving spatial distribution of residents in colony biofilms: an interdisciplinary perspective. Open Biology 12:220194. https://royalsocietypublishing.org/doi/10.1098/rsob.220194.

Hallatschek, O., and D. R. Nelson, 2010. Life at the front of an expanding population. Evolution 64:193–206. https://academic.oup.com/evolut/article/64/1/193/6854125.

Branda, S. S., Ã. Vik, L. Friedman, and R. Kolter, 2005. Biofilms: the matrix revisited. Trends in Microbiology 13:20–26. https://linkinghub.elsevier.com/retrieve/pii/S0966842X04002604.

Hallatschek, O., P. Hersen, S. Ramanathan, and D. R. Nelson, 2007. Genetic drift at expanding frontiers promotes gene segregation. Proceedings of the National Academy of Sciences 104:19926–19930. https://pnas.org/doi/full/10.1073/pnas.0710150104.

Korolev, K. S., J. B. Xavier, D. R. Nelson, and K. R. Foster, 2011. A Quantitative Test of Population Genetics Using Spatiogenetic Patterns in Bacterial Colonies. The American Naturalist 178:538–552. https://www.journals.uchicago.edu/doi/10.1086/661897.

Excoffier, L., and N. Ray, 2008. Surfing during population expansions promotes genetic revolutions and structuration. Trends in Ecology & Evolution 23:347–351. https://linkinghub.elsevier.com/retrieve/pii/S0169534708001675.

Jauffred, L., R. Munk Vejborg, K. S. Korolev, S. Brown, and L. B. Oddershede, 2017. Chirality in microbial biofilms is mediated by close interactions between the cell surface and the substratum. The ISME Journal 11:1688–1701. https://www.nature.com/articles/ismej201719.

Booth, S. C., and S. A. Rice, 2020. Influence of interspecies interactions on the spatial organization of dual species bacterial communities. Biofilm 2:100035. https://linkinghub.elsevier.com/retrieve/pii/S2590207520300174.

Smith, W. P. J., Y. Davit, J. M. Osborne, W. D. Kim, K. R. Foster, and J. M. Pitt-Francis, 2017. Cell morphology drives spatial patterning in microbial communities. Proceedings of the National Academy of Sciences 114:E280–E286. http://www.pnas.org/lookup/doi/10.1073/pnas.1613007114 10.1073/pnas.1613007114.

van Gestel, J., F. J. Weissing, O. P. Kuipers, and Ã. T. Kovács, 2014. Density of founder cells affects spatial pattern formation and cooperation in Bacillus subtilis biofilms. The ISME Journal 8:2069–2079. https://www.nature.com/articles/ismej201452.

Blanchard, A. E., and T. Lu, 2015. Bacterial social interactions drive the emergence of differential spatial colony structures. BMC Systems Biology 9:59. https://bmcsystbiol.biomedcentral.com/articles/10.1186/s12918-015-0188-5.

Borenstein, D. B., P. Ringel, M. Basler, and N. S. Wingreen, 2015. Established Microbial Colonies Can Survive Type VI Secretion Assault. PLOS Computational Biology 11:e1004520. https://dx.plos.org/10.1371/journal.pcbi.1004520.

Bottery, M. J., I. Passaris, C. Dytham, A. J. Wood, and M. W. van der Woude, 2019. Spatial Organization of Expanding Bacterial Colonies Is Affected by Contact-Dependent Growth Inhibition. Current Biology 29:3622–3634. https://linkinghub.elsevier.com/retrieve/pii/S0960982219311558.

Lee, H., J. Gore, and K. S. Korolev, 2022. Slow expanders invade by forming dented fronts in microbial colonies. Proceedings of the National Academy of Sciences 119:e2108653119. https://pnas.org/doi/full/10.1073/pnas.2108653119.

Eigentler, L., M. Kalamara, G. Ball, C. E. MacPhee, N. R. Stanley-Wall, and F. A. Davidson, 2022. Founder cell configuration drives competitive outcome within colony biofilms. The ISME Journal 16:1512–1522. https://www.nature.com/articles/s41396-022-01198-8.

Korolev, K. S., M. Avlund, O. Hallatschek, and D. R. Nelson, 2010. Genetic demixing and evolution in linear stepping stone models. Reviews of Modern Physics 82:1691–1718. https://link.aps.org/doi/10.1103/RevModPhys.82.1691.

Fusco, D., M. Gralka, J. Kayser, A. Anderson, and O. Hallatschek, 2016. Excess of mutational jackpot events in expanding populations revealed by spatial LuriaâĂŞDelbrück experiments. Nature Communications 7:12760. https://www.nature.com/articles/ncomms12760.

Jung, Y.-G., J. Choi, S.-K. Kim, J.-H. Lee, and S. Kwon, 2015. Embedded Biofilm, a New Biofilm Model Based on the Embedded Growth of Bacteria. Applied and Environmental Microbiology 81:211–219. https://journals.asm.org/doi/10.1128/AEM.02311-14.

Strathmann, M., T. Griebe, and H. C. Flemming, 2000. Artificial biofilm model-A useful tool for biofilm research. Applied Microbiology and Biotechnology 54:231–237. https://link.springer.com/content/pdf/10.1007/s002530000370.pdf.

Burmølle, M., K. Johnsen, W. A. Al-Soud, L. H. Hansen, and S. J. Sørensen, 2009. The presence of embedded bacterial pure cultures in agar plates stimulate the culturability of soil bacteria. Journal of Microbiological Methods 79:166–173. 10.1016/j.mimet.2009.08.006.

Zhang, Q., J. Li, J. Nijjer, H. Lu, M. Kothari, R. Alert, T. Cohen, and J. Yan, 2021. Morphogenesis and cell ordering in confined bacterial biofilms. Proceedings of the National Academy of Sciences 118. https://pnas.org/doi/full/10.1073/pnas.2107107118.

Cordero, M., N. Mitarai, and L. Jauffred, 2023. Motility mediates satellite formation in confined biofilms. The ISME Journal 17:1819–1827. https://www.nature.com/articles/s41396-023-01494-x.

Müller, J., A. C. Jäkel, J. Richter, M. Eder, E. Falgenhauer, and F. C. Simmel, 2022. Bacterial Growth, Communication, and Guided Chemotaxis in 3D-Bioprinted Hydrogel Environments. ACS Applied Materials & Interfaces 14:15871–15880. https://pubs.acs.org/doi/10.1021/acsami.1c20836.

Eriksen, R. S., S. L. Svenningsen, K. Sneppen, and N. Mitarai, 2018. A growing microcolony can survive and support persistent propagation of virulent phages. Proceedings of the National Academy of Sciences 115:337–342. http://www.pnas.org/lookup/doi/10.1073/pnas.1708954115.

Ben-Jacob, E., D. S. Coffey, and H. Levine, 2012. Bacterial survival strategies suggest rethinking cancer cooperativity. Trends in Microbiology 20:403–410. 10.1016/j.tim.2012.06.001.

Buffi, N., D. Merulla, J. Beutier, F. Barbaud, S. Beggah, H. van Lintel, P. Renaud, and J. Roelof van der Meer, 2011. Development of a microfluidics biosensor for agarose-bead immobilized Escherichia coli bioreporter cells for arsenite detection in aqueous samples. Lab on a Chip 11:2369. http://xlink.rsc.org/?DOI=c1lc20274j.

Kuennen, E. W., and C. Y. Wang, 2008. Off-lattice radial Eden cluster growth in two and three dimensions. Journal of Statistical Mechanics: Theory and Experiment 2008:P05014. https://iopscience.iop.org/article/10.1088/1742-5468/2008/05/P05014.

Shao, X., A. Mugler, J. Kim, H. J. Jeong, B. R. Levin, and I. Nemenman, 2017. Growth of bacteria in 3-d colonies. PLOS Computational Biology 13:e1005679. https://dx.plos.org/10.1371/journal.pcbi.1005679.

Lavrentovich, M. O., and D. R. Nelson, 2015. Survival probabilities at spherical frontiers. Theoretical Population Biology 102:26–39. https://linkinghub.elsevier.com/retrieve/pii/S0040580915000210.

Jullien, R., and R. Botet, 1985. Surface Thickness in the Eden Model. Physical Review Letters 54:2055–2055. https://link.aps.org/doi/10.1103/PhysRevLett.54.2055.

Bredenoord, A. L., H. Clevers, and J. A. Knoblich, 2017. Human tissues in a dish: The research and ethical implications of organoid technology. Science 355:eaaf9414. https://www.science.org/doi/10.1126/science.aaf9414.

Weiswald, L. B., D. Bellet, and V. Dangles-Marie, 2015. Spherical cancer models in tumor biology. Neoplasia (New York, N.Y.) 17:1–15. 10.1016/j.neo.2014.12.004.

Costa, E. C., A. F. Moreira, D. de Melo-Diogo, V. M. Gaspar, M. P. Carvalho, and I. J. Correia, 2016. 3D tumor spheroids: an overview on the tools and techniques used for their analysis. Biotechnology Advances 34:1427–1441. 10.1016/j.biotechadv.2016.11.002.

Tomer, R., K. Khairy, F. Amat, and P. J. Keller, 2012. Quantitative high-speed imaging of entire developing embryos with simultaneous multiview light-sheet microscopy. Nature Methods 9:755–63. http://www.ncbi.nlm.nih.gov/pubmed/22660741.

Boyer, D., W. Mather, O. Mondragón-Palomino, S. Orozco-Fuentes, T. Danino, J. Hasty, and L. S. Tsimring, 2011. Buckling instability in ordered bacterial colonies. Physical Biology 8:026008. https://iopscience.iop.org/article/10.1088/1478-3975/8/2/026008.

Rudge, T. J., F. Federici, P. J. Steiner, A. Kan, and J. Haseloff, 2013. Cell Polarity-Driven Instability Generates Self-Organized, Fractal Patterning of Cell Layers. ACS Synthetic Biology 2:705–714. https://pubs.acs.org/doi/10.1021/sb400030p.

Kan, A., I. Del Valle, T. Rudge, F. Federici, and J. Haseloff, 2018. Intercellular adhesion promotes clonal mixing in growing bacterial populations. Journal of the Royal Society Interface 15:20180406. https://royalsocietypublishing.org/doi/10.1098/rsif.2018.0406.

Tuson, H. H., G. K. Auer, L. D. Renner, M. Hasebe, C. Tropini, M. Salick, W. C. Crone, A. Gopinathan, K. C. Huang, and D. B. Weibel, 2012. Measuring the stiffness of bacterial cells from growth rates in hydrogels of tunable elasticity. Molecular Microbiology 84:874–891. https://onlinelibrary.wiley.com/doi/10.1111/j.1365-2958.2012.08063.x.

Lavrentovich, M. O., J. H. Koschwanez, and D. R. Nelson, 2013. Nutrient shielding in clusters of cells. Physical Review E 87:062703. https://link.aps.org/doi/10.1103/PhysRevE.87.062703.

Martínez-Calvo, A., T. Bhattacharjee, R. K. Bay, H. N. Luu, A. M. Hancock, N. S. Wingreen, and S. S. Datta, 2022. Morphological instability and roughening of growing 3D bacterial colonies. Proceedings of the National Academy of Sciences 119:819–825. https://pnas.org/doi/10.1073/pnas.2208019119.

Ohgiwari, M., M. Matsushita, and T. Matsuyama, 1992. Morphological Changes in Growth Phenomena of Bacterial Colony Patterns. Journal of the Physical Society of Japan 61:816–822. https://journals.jps.jp/doi/10.1143/JPSJ.61.816.

Studier, F. W., P. Daegelen, R. E. Lenski, S. Maslov, and J. F. Kim, 2009. Understanding the Differences between Genome Sequences of Escherichia coli B Strains REL606 and BL21(DE3) and Comparison of the E. coli B and K-12 Genomes. Journal of Molecular Biology 394:653–680. https://linkinghub.elsevier.com/retrieve/pii/S0022283609011383.

Lavrentovich, M. O., K. S. Korolev, and D. R. Nelson, 2013. Radial Domany-Kinzel models with mutation and selection. Physical Review E 87:012103. https://link.aps.org/doi/10.1103/PhysRevE.87.012103.

Castillo, C. E., and M. O. Lavrentovich, 2020. Shape of population interfaces as an indicator of mutational instability in coexisting cell populations. Physical Biology 17:066002. https://iopscience.iop.org/article/10.1088/1478-3975/abb2dd.

Dornic, I., H. Chaté, J. Chave, and H. Hinrichsen, 2001. Critical Coarsening without Surface Tension: The Universality Class of the Voter Model. Physical Review Letters 87:045701. https://link.aps.org/doi/10.1103/PhysRevLett.87.045701.

Kuhr, J.-T., M. Leisner, and E. Frey, 2011. Range expansion with mutation and selection: dynamical phase transition in a two-species Eden model. New Journal of Physics 13:113013. https://iopscience.iop.org/article/10.1088/1367-2630/13/11/113013.

Friedrich, J., C. Seidel, R. Ebner, and L. A. Kunz-Schughart, 2009. Spheroid-based drug screen: considerations and practical approach. Nature Protocols 4:309–324. http://www.nature.com/doifinder/10.1038/nprot.2008.226.

Dal Co, A., S. van Vliet, D. J. Kiviet, S. Schlegel, and M. Ackermann, 2020. Short-range interactions govern the dynamics and functions of microbial communities. Nature Ecology & Evolution 4:366–375. https://www.nature.com/articles/s41559-019-1080-2.

Borer, B., D. Ciccarese, D. Johnson, and D. Or, 2020. Spatial organization in microbial range expansion emerges from trophic dependencies and successful lineages. Communications Biology 3:685. https://www.nature.com/articles/s42003-020-01409-y.

Celik Ozgen, V., W. Kong, A. E. Blanchard, F. Liu, and T. Lu, 2018. Spatial interference scale as a determinant of microbial range expansion. Science Advances 4. https://www.science.org/doi/10.1126/sciadv.aau0695.

Monds, R. D., T. K. Lee, A. Colavin, T. Ursell, S. Quan, T. F. Cooper, and K. C. Huang, 2014. Systematic Perturbation of Cytoskeletal Function Reveals a Linear Scaling Relationship between Cell Geometry and Fitness. Cell Reports 9:1528–1537. 10.1016/j.celrep.2014.10.040.

Elbing, K., and R. Brent, 2002. Media Preparation and Bacteriological Tools. Current Protocols in Molecular Biology 59:1–1. https://onlinelibrary.wiley.com/doi/10.1002/0471142727.mb0101s59.

Mao, B., A. Bentaleb, F. Louerat, T. Divoux, and P. Snabre, 2017. Heat-induced aging of agar solutions: Impact on the structural and mechanical properties of agar gels. Food Hydrocolloids 64:59–69. https://linkinghub.elsevier.com/retrieve/pii/S0268005X16305793.

Sønderholm, M., K. N. Kragh, K. Koren, T. H. Jakobsen, S. E. Darch, M. Alhede, P. Ã. Jensen, M. Whiteley, M. Kühl, and T. Bjarnsholt, 2017. Pseudomonas aeruginosa Aggregate Formation in an Alginate Bead Model System Exhibits <i>In Vivo</i> -Like Characteristics. Applied and Environmental Microbiology 83:1–15. https://journals.asm.org/doi/10.1128/AEM.00113-17.

Kvich, L., B. Fritz, S. Crone, K. N. Kragh, M. Kolpen, M. Sønderholm, M. Andersson, A. Koch, P. Ã. Jensen, and T. Bjarnsholt, 2019. Oxygen Restriction Generates Difficult-to-Culture P. aeruginosa. Frontiers in Microbiology 10:1992. https://www.frontiersin.org/article/10.3389/fmicb.2019.01992/full.

Schmid, B., J. Schindelin, A. Cardona, M. Longair, and M. Heisenberg, 2010. A high-level 3D visualization API for Java and ImageJ. BMC Bioinformatics 11:274. https://bmcbioinformatics.biomedcentral.com/articles/10.1186/1471-2105-11-274.

Preibisch, S., S. Saalfeld, and P. Tomancak, 2009. Globally optimal stitching of tiled 3D microscopic image acquisitions. Bioinformatics 25:1463–1465. https://academic.oup.com/bioinformatics/article/25/11/1463/332497.

Hartmann, R., H. Jeckel, E. Jelli, P. K. Singh, S. Vaidya, M. Bayer, D. K. H. Rode, L. Vidakovic, F. Díaz-Pascual, J. C. N. Fong, A. Dragoš, O. Lamprecht, J. G. Thöming, N. Netter, S. Häussler, C. D. Nadell, V. Sourjik, Ã. T. Kovács, F. H. Yildiz, and K. Drescher, 2021. Quantitative image analysis of microbial communities with BiofilmQ. Nature Microbiology 6:151–156. https://www.nature.com/articles/s41564-020-00817-4.

Schindelin, J., I. Arganda-Carreras, E. Frise, V. Kaynig, M. Longair, T. Pietzsch, S. Preibisch, C. Rueden, S. Saalfeld, B. Schmid, J.-Y. Tinevez, D. J. White, V. Hartenstein, K. Eliceiri, P. Tomancak, and A. Cardona, 2012. Fiji: an open-source platform for biological-image analysis. Nature Methods 9:676–682. https://www.nature.com/articles/nmeth.2019.

Ollion, J., J. Cochennec, F. Loll, C. Escudé, and T. Boudier, 2013. TANGO: a generic tool for high-throughput 3D image analysis for studying nuclear organization. Bioinformatics 29:1840–1841. https://academic.oup.com/bioinformatics/article/29/14/1840/231770.

Sanderson, C., and R. Curtin, 2016. Armadillo: a template-based C++ library for linear algebra. The Journal of Open Source Software 1:26. http://joss.theoj.org/papers/10.21105/joss.00026.

