## Supplementary figures and images for "Genetic mixing and demixing on expanding spherical frontiers"

### Movie 1

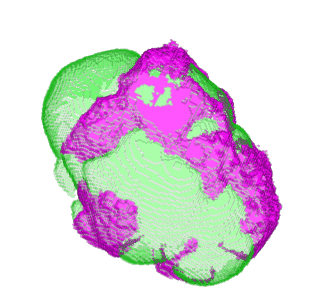

### Movie 2

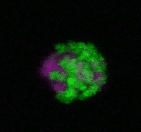

### Movie 3

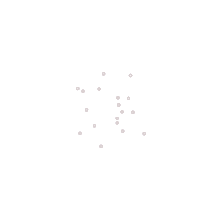

### Movie 4

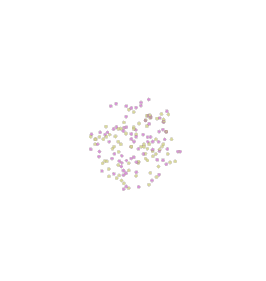

### Movie 5

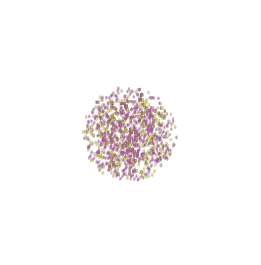

### Movie 6

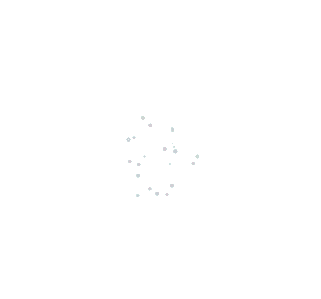

### Movie 7

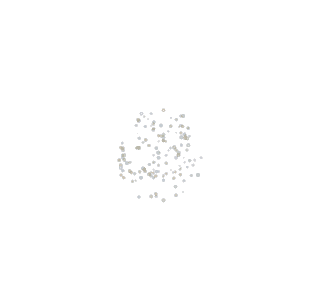

### Movie 8

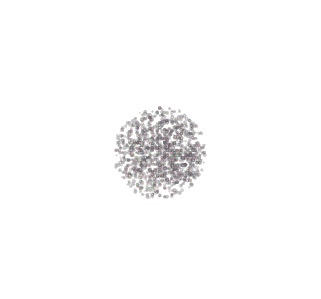
