## Supplementary material for "Genetic mixing and demixing on expanding spherical frontiers": SI

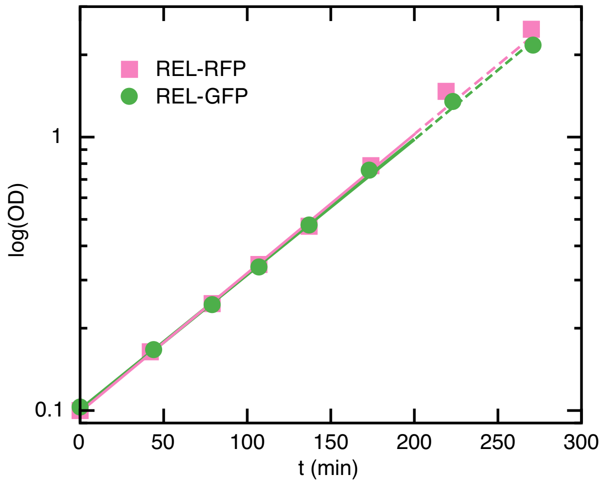


**Figure S1:** Growth rates for REL *E. coli*. Optical density, OD, verus time, $t$, on a semi-logarithmic scale in base 2 of REL-RFP (magenta) and REL-GFP (green). Linear fits (full lines) gives the doubling times: (59±1) min and (60±1) min for REL-GFP and REL-RFP, respectively. The two last points were excluded from the fit since OD> 1 is often out of exponential phase (1, 2).

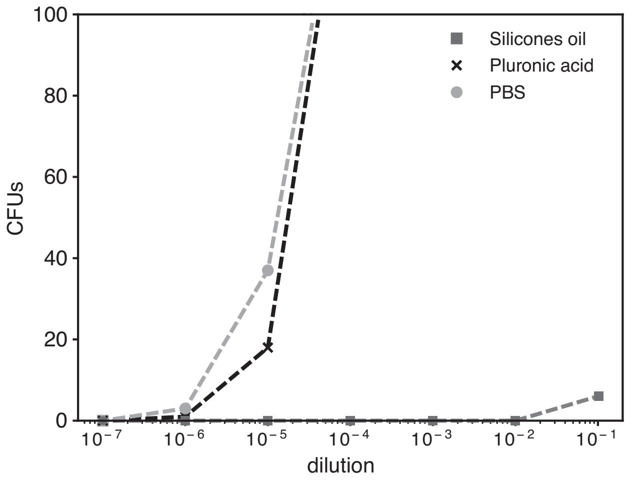


**Figure S2:** Toxicity of silicone oil (dark grey squares), pluronic acid (black crosses) and the control PBS (light grey circles). CFU counts versus dilution (10× - $10^7$× diluted) on semi-logarithmic scales.

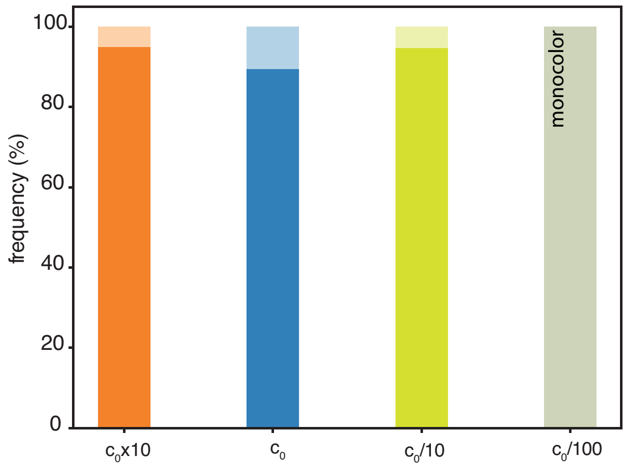


**Figure S3:** Frequency of two-colored (full color) and monocolor (opaque) colonies derived from inoculation beads with concentration: $c_0 \times 10$ (N=20), $c_0$ (N=19), $c_0/10$ (N=19), and $c_0/100$ (N=5).

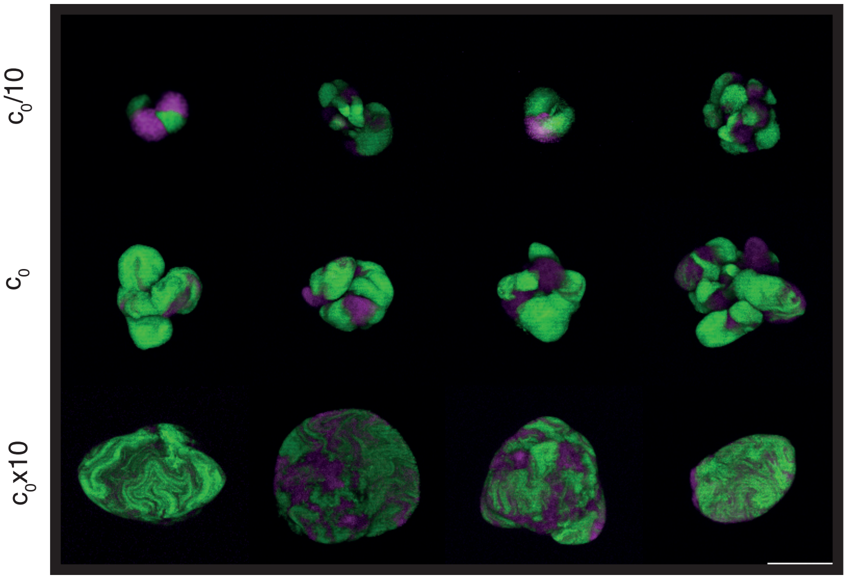


**Figure S4:** Maximum-intensity z-projected examples of colonies derived from inoculation beads with concentration: $c_0/10$, $c_0$, $c_0 \times 10$. The scale bar corresponds to 100 $\mu$m

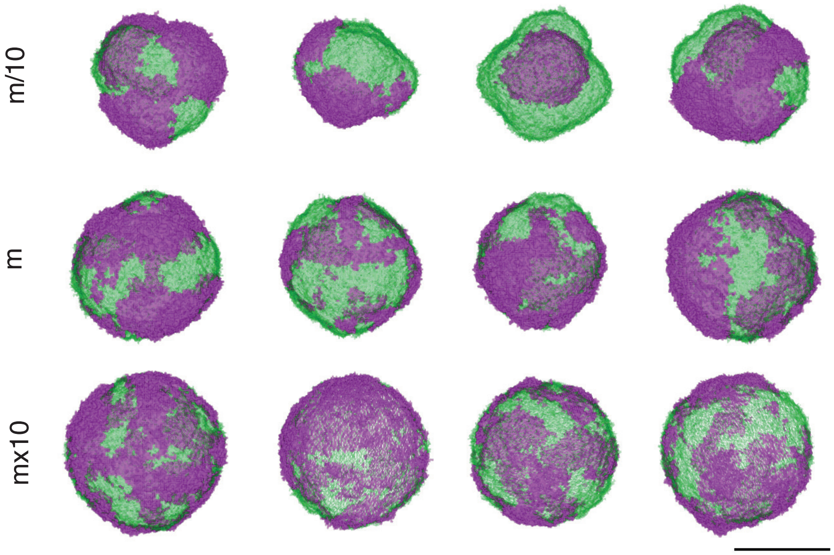


**Figure S5:** Examples of two-species *in silico* colonies derived from seeding spheres with concentration: $m/10$, $m$, $m \times 10$. The scale bar corresponds to 50 lattice sites.

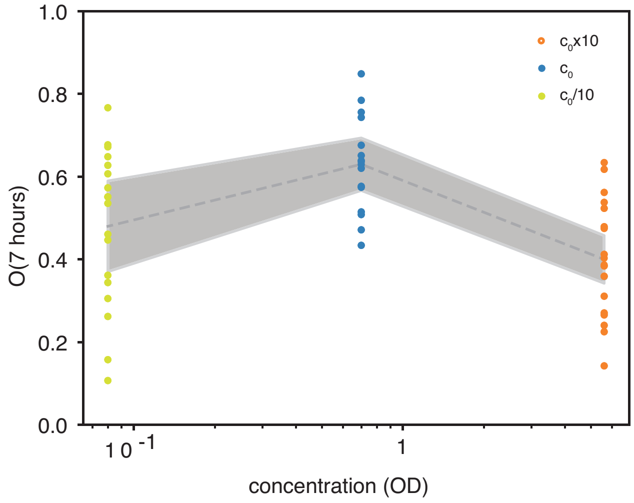


**Figure S6:** Fraction of GFP-expressing sub-population on the colonies' surface, $O(t)$, at $t = 7$ hours as defined in Equation 3, for $c_0/10$ (N=18), $c_0$ (N=17), and $c_0 \times 10$ (N=18). The shaded region corresponds to ±SEM.

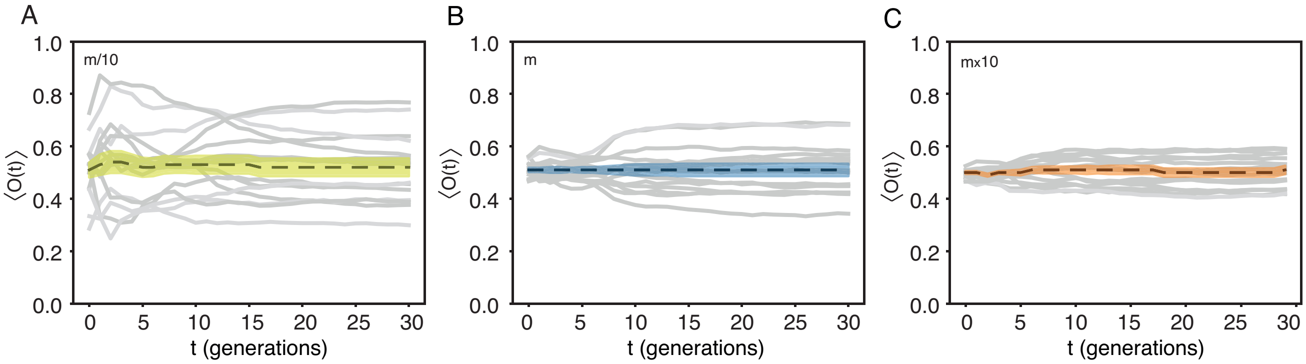


**Figure S7:** Fraction of one sub-population on the colonies' surface, $O(t)$, versus time, $t$, as defined in Equation 2, for *in silico* colonies at 3 different initial concentrations: $m/10$ seeds (A), $m$ seeds (B), and $m \times 10$ seeds (C). The punctuated line is the mean of all colonies and the shaded area signify ±SEM.

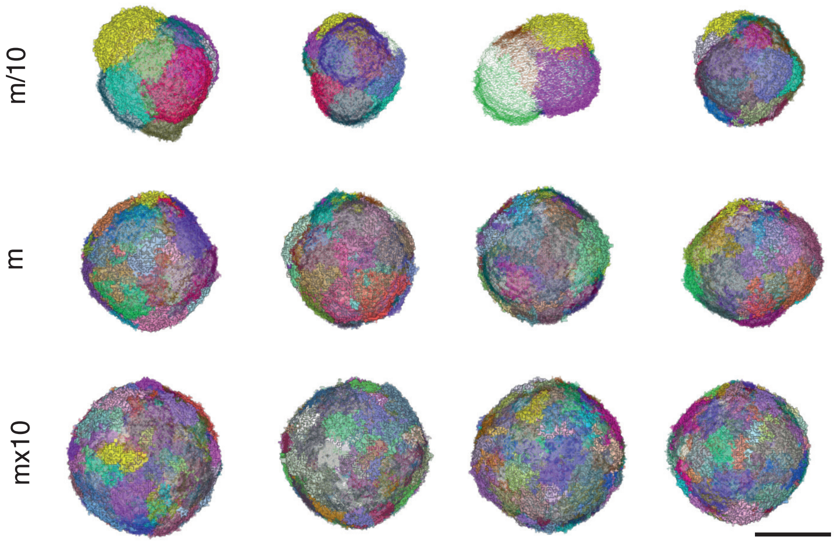


**Figure S8:** Examples of multi-species *in silico* colonies derived from seeding spheres with concentration: $m/10$, $m$, $m \times 10$. The scale bar corresponds to 50 lattice sites.

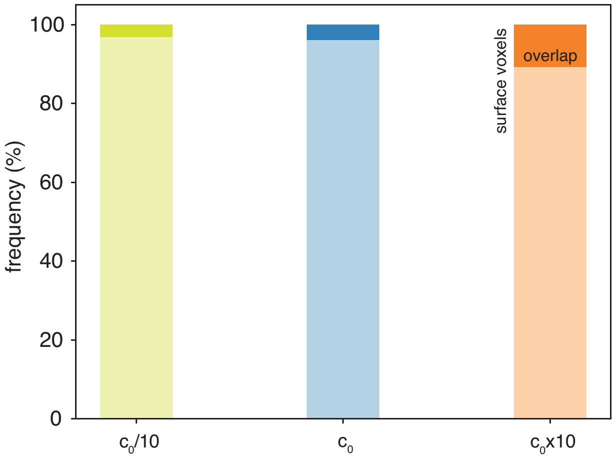


**Figure S9:** Frequency of surface voxels assigned both colours (overlap) from all (surface voxels), i.e., full + opaque region. The frequency is the average values (3.1±0.5)%, (3.8±0.4)%, and (10.7±1.2)% for $c_0/10$ (N=18), $c_0$ (N=17), and $c_0 \times 10$ (N=19), respectively.

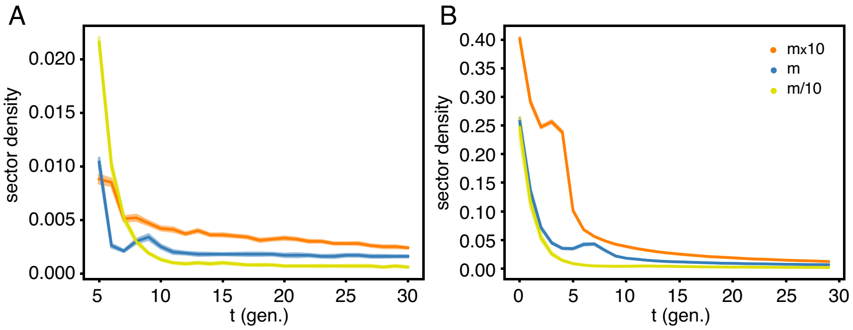


**Figure S10:** Ensemble-averaged sector density for two-species (A) and multi-species (B) *in silico* colonies at 3 different initial concentrations: $m/10$ seeds (yellow), $m$ seeds (blue), and $m \times 10$ seeds (orange). The fraction of sectors for the $n$-th colony is $\sigma^{(n)}(t)/A^{(n)}(t)$ versus time, $t$. The punctuated line is the average over all colonies and the shaded area signify ±SEM.

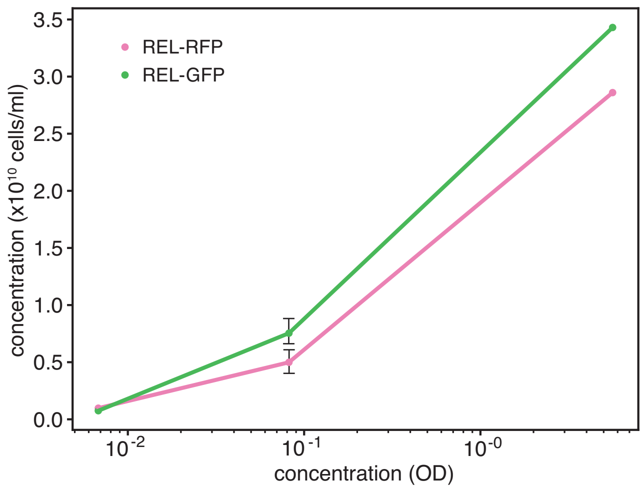


**Figure S11:** Number of CFUs pr. volume, REL-RFP (magenta) and REL-GFP (green), versus density of cells (OD) in the final inoculation beads: $c_0/100$, $c_0/10$, $c_0$. Error bars signifies one SD (N=2).

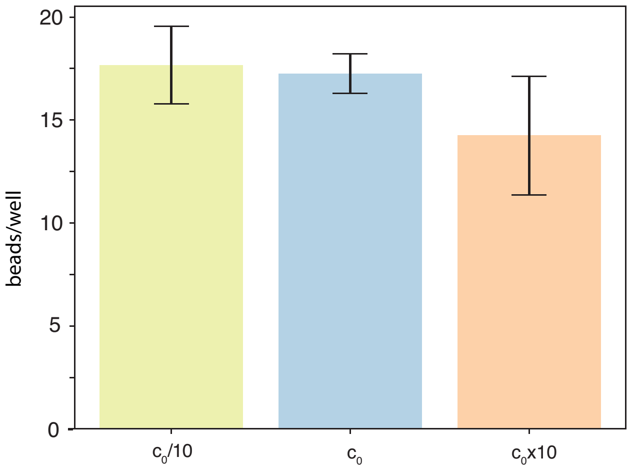


**Figure S12:** Number of colonies per well. The average number of colonies per well for the concentrations: $c_0 \times 10$ (N=6), $c_0$ (N=4), and $c_0/10$ (N=4). The error bars signifies one SEM.

### MOVIE LEGENDS

#### Movie 1

360 degrees recording of an example mask, $M$, obtained 3D image originating from a colony incubated for 7 hours from an inoculation bead of concentration $c_0$. All voxels were assigned one (and only one) colour.

#### Movie 2

Maximum intensity z-projections time-lapse (frame rate is 4/hour) starting after 5 hours of incubation. The colony originates from an inoculation bead of concentration $c_0$.

#### Movie 3

Time-lapse (frame rate is 1/generations) of an two-species *in silico* colony for 29 generations starting from a seeding concentration of $m/10$.

#### Movie 4

Time-lapse (frame rate is 1/generations) of an two-species *in silico* colony for 29 generations starting from a seeding concentration of $m$.

#### Movie 5

Time-lapse (frame rate is 1/generations) of an two-species *in silico* colony for 29 generations starting from a seeding concentration of $m \times 10$.

#### Movie 6

Time-lapse (frame rate is 1/generations) of an multi-species *in silico* colony for 29 generations starting from a seeding concentration of $m/10$.

#### Movie 7

Time-lapse (frame rate is 1/generations) of an multi-species *in silico* colony for 29 generations starting from a seeding concentration of $m$.

#### Movie 8

Time-lapse (frame rate is 1/generations) of an multi-species *in silico* colony for 29 generations starting from a seeding concentration of $m \times 10$.

### REFERENCES

1. Kobayashi, A., H. Hirakawa, T. Hirata, K. Nishino, and A. Yamaguchi, 2006. Growth phase-dependent expression of drug exporters in Escherichia coli and its contribution to drug tolerance. *Journal of Bacteriology* 188.

2. Tsuchiya, K., Y. Y. Cao, M. Kurokawa, K. Ashino, T. Yomo, and B. W. Ying, 2018. A decay effect of the growth rate associated with genome reduction in Escherichia coli. *BMC Microbiology* 18.
